# Carbon Cybernetics Array: a miniaturized carbon-based microelectrode array for intracortical recording

**DOI:** 10.64898/2026.01.06.696090

**Authors:** Simon Higham, Young Jun Jung, Sorel E. De Leon, Wei Tong, Shun Long Cyril Au, Yidan Shang, Han Wang, Mauhak Kalra, Andrew Morokoff, Matias I. Maturana, Rob Hilkes, Michael R. Ibbotson, David J. Garrett, Steven Prawer

**Affiliations:** School of Physics, The University of Melbourne, Melbourne, Victoria, Australia; Department of Biomedical Engineering, RMIT University, Melbourne, Victoria, Australia; Department of Biomedical Engineering, The University of Melbourne, Melbourne, Victoria, Australia; The Graeme Clark Institute, The University of Melbourne, Melbourne, Victoria, Australia; Carbon Cybernetics Canada Ltd, Ottawa, Canada; School of Mechanical and Automotive Engineering, Shanghai University of Engineering Science, Shanghai, China; Department of Surgery, The University of Melbourne, Melbourne, Victoria, Australia

**Keywords:** carbon fiber, diamond, neural recording, neural interface, implantable

## Abstract

Implantable brain-machine interfaces, capable of capturing high-resolution neural signals, hold great promise for treating neurological conditions and advancing neuroscience research. As the field moves toward chronic, fully implantable systems, ensuring the long-term safety and stability of the microelectrode arrays, the core component of these interfaces, remains a significant challenge. Hermetic sealing, which prevents moisture and contaminants from entering the device, is a critical yet often overlooked factor in array designs. This study reports the Carbon Cybernetics Array, a hermetically sealed, carbon-based microelectrode array for neural interfacing. Carbon fiber electrodes are anchored on a diamond substrate via a hermetic feedthrough structure, with hermeticity validated through helium leak testing. These ultrathin and flexible carbon fibers require minimal insertion force, cause little tissue damage after six months of implantation in rats and enable single-unit neural recordings. Furthermore, a low-cost, manually controllable insertion toolset is developed for precise array implantation in the sheep brain through a small burr hole, demonstrating a step toward clinical translation. Finally, the scalability of the fabrication approach is exemplified by arrays with over 1000 fibers. Together, this hermetically sealed feedthrough structure will lay the foundation for a future fully implantable device with wireless capabilities.

## 1. INTRODUCTION

Brain-machine interfaces (BMIs) provide a direct communication pathway between the brain and the external world through neural stimulation and recording. Recent years have seen exciting advances in the use of BMIs to restore lost functions in individuals with severe neurological disorders, promoting greater independence^1^. Intracortical BMIs, which involve surgically implanting sensors into the cortex, allow for the assessment of information-rich single unit activity and local field potentials without signal degradation caused by spatial averaging or bone filtering, offering higher-resolution communication compared to less invasive approaches such as electroencephalogram (EEG) and electrocorticography (ECoG)^2^. Successful applications include using neural signals to control computer cursors^3^, manipulate robotic arms^4^, and restore rapid communication through accurate decoding of handwriting^5^ and speech^6^.

While the development of algorithms to analyze neural signals has advanced rapidly, the availability of implantable devices for clinical applications remains limited. This disparity underscores the challenge of translating fabrication techniques into clinically-viable, long-term solutions for implantable neural interfaces. The Utah array (Neuroport, Blackrock Microsystems Inc.), which is composed of 96 silicon-based probes^7^, remains one of the most used intracortical microelectrode arrays for use in human trials, despite Neuralink Corp. launching its clinical trials using 1024 electrodes distributed across 64 shanks^8^. Concerns regarding the longevity and stability of Utah arrays persist, particularly due to the adverse tissue response around the electrodes after chronic implantation, as observed in both animal^9^ and clinical studies^7, 10^. To address this issue, researchers have explored using flexible, softer materials that offer better compatibility with neural tissue, such as the Neuralink^11^ and the nanoelectronic thread (NET) electrodes^12^. These designs also differ from the Utah array in structure: while the Utah array consists of multiple single-electrode shanks and provides lateral information from the cortex, Neuralink and NET electrodes use shanks that each contain multiple electrodes for communicating at different depths within the cortex. Consequently, existing applications and data analysis algorithms developed for Utah arrays may not seamlessly adapt to these new configurations. While alternative arrays with similar structures to Utah arrays have merged, like the CMOS microwire array^13^, the Argo system^14^ by Paradromics Inc. and the CMU array^15^, the prevalent use of rigid materials such as metal and silicon raises questions about their long-term biocompatibility and stability.

Presently, the trajectory of the field is steering towards the development of fully implantable devices^16–18^. These innovative designs emphasize miniaturization and wireless capabilities, in contrast to the tethered design of the NeuroPort system used with the Utah arrays^7^. The benefits of such advances include less invasive surgical procedures for easier implantation (and explantation) without brain injury, enhanced patient experience and clinical outcomes, and improved device functionality^16^.

Hermeticity is a crucial factor in the construction of electrode arrays for fully implantable devices^16, 19^. The environment to which these implanted arrays are exposed is both chemically diverse and aggressive. Reactive oxygen species produced by the immune response can attack and degrade the implant, leading to the loss of hermeticity. This can result in moisture and other contaminants entering the package, which can lead to corrosion of the microelectronics and short-circuits. Conversely, potential leaching of non-biocompatible materials can cause cell death and exacerbate the immune response, consequently limiting the device’s operational lifetime.

Carbon fibers, characterized by their ultra-small diameter (<10 µm) and flexibility, have emerged as promising electrode materials for neural interfaces due to their potential to reduce glial scarring, making them suitable for long-term applications^20–23^. With appropriate surface coatings, these carbon fiber electrodes have been used for single-unit recording^20–25^, electrical stimulation^24, 25^ and electrochemical sensing^24, 26^. While multiple research groups have successfully integrated these carbon fibers into array configurations, many are restrained by a large electrode pitch (larger than 150 µm) and predominantly feature 2-column electrode layouts^21, 26^. Novel fabrication techniques have emerged for fabricating arrays with just 20 µm pitch between recording electrodes^22^. However, these pioneering designs typically anchor the carbon fibers to substrates like conventional printed circuit boards^21, 26^, 3D-printed plastic blocks^23^, or silicon substrates^22^. To date, none of these configurations facilitate hermetic feedthrough structures, a critical feature for the realisation of future fully implantable devices for long-term applications. Furthermore, all the current designs use silver paste to connect carbon fibers to external electronics, which is known to be toxic to humans.

In this paper, we introduce the Carbon Cybernetics array: a miniaturized, carbon-based microelectrode array for intracortical recording (**Figure 1**). Using advanced diamond technology, we have assembled carbon fibers into a configuration mirroring the Utah array structure, with a compact electrode pitch of as small as 100 µm (**Figure 1A**). These fibers are secured to a 2.5 mm diameter insulating diamond substrate. The inherent biocompatible and bioinert nature of diamond enhance both the longevity and safety of our design. The electrical connection to carbon fibers is achieved by crafting a microcircuit directly on the reverse side of the diamond substrate using laser micro-machining and brazing (**Figure 1B**). We show evidence of the hermeticity of this feedthrough structure via helium leakage tests, making it suitable for medical implant applications. We also demonstrate the long-term safety of the electrodes in rat cortex, showing a minimal tissue response after six months of implantation. Moreover, the capability of these arrays for *in vivo* neural recording at single-neuron resolution is validated. To facilitate translation to large animals and human applications, we developed a low-cost, manually controllable 3D-printed inserter toolset. This toolset enables precise implantation of the arrays into the sheep’s brain via a small burr hole, removing the need for large and expensive insertion robotics and extensive, risky craniotomies. We further demonstrate the scalability of our fabrication methods, with arrays of over 1000 channels fabricated. Our routing method on diamond paves the way for the subsequent integration of an application-specific integrated circuit (ASIC)^27^, laying the foundation for a compact, fully implantable wireless neural interface system. Such a miniaturised system promises to bring significant benefit to the neuroscience community and offer innovative treatments for diverse neurological disorders.

**Figure 1.**
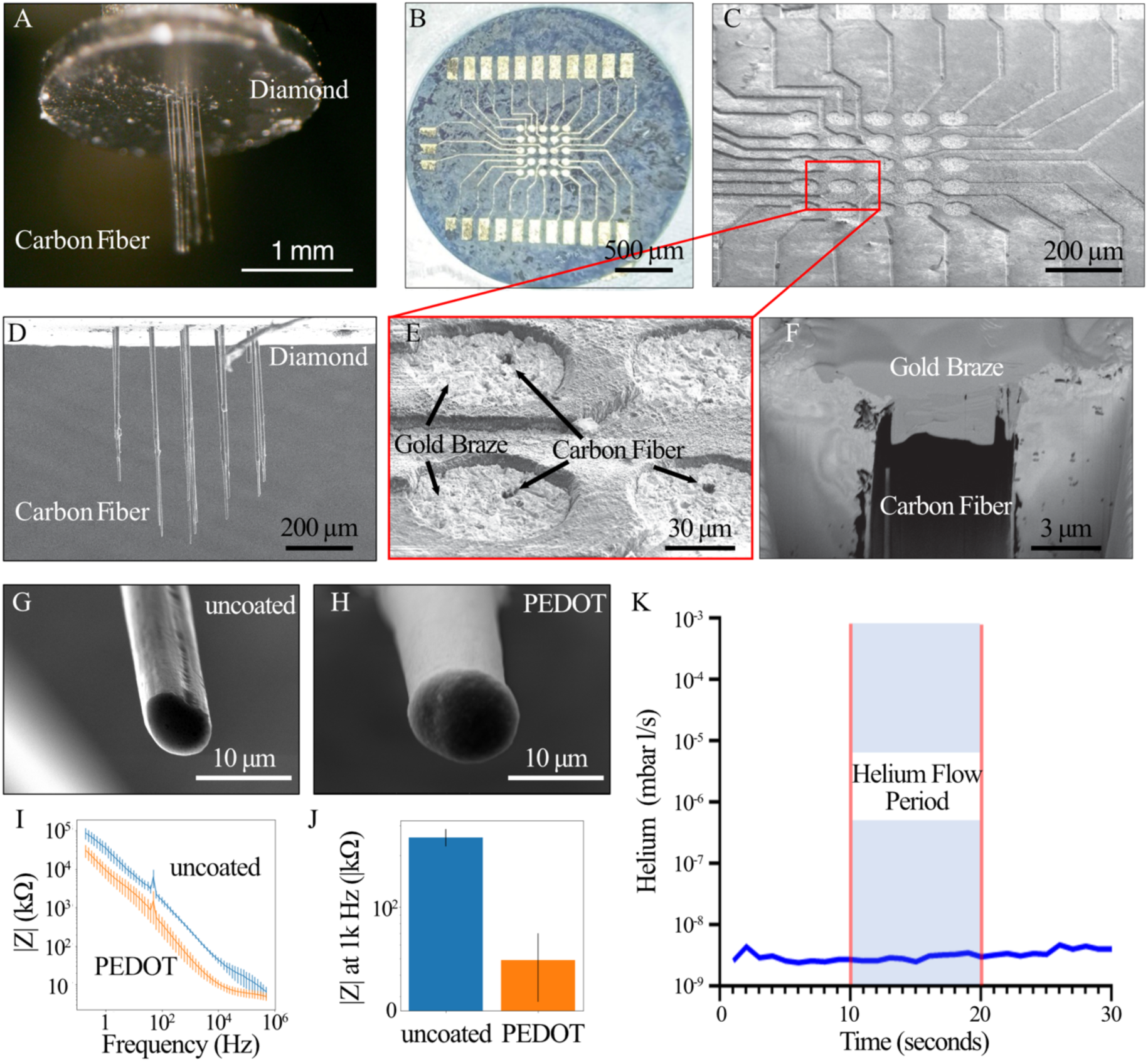
The Carbon Cybernetics Array Fabrication and Hermeticity Test. (A) A macro photograph showing an array comprising 25 carbon fiber electrodes anchored on a diamond substrate. (B) A microscope image of the diamond circuit board for a 25-carbon fiber array. This circuit board is designed to bridge the carbon fiber electrodes with other electronic components. The central 5 × 5 grid are feedthrough holes for carbon fibers, which are connected to the peripheral pads on the diamond substrate for other electronic interfacing. (C) SEM image of the diamond circuit board, highlighting the complete filling of the braze of the feedthrough holes and the routing traces. (D) SEM image of a carbon fiber array with 25 fibers of different lengths. (E) Close-up view of (C) showing visible carbon fibers on the gold braze surface within the feedthrough holes. (F) SEM cross-section around a carbon fiber, prepared via FIB, demonstrating the intimate contact between the carbon fiber and gold braze. (G) SEM image of an uncoated carbon fiber tip. (H) SEM image after PEDOT:PSS coating. (I) Electrochemical impedance spectroscopy before and after PEDOT: PSS application. (J) Change of electrode impedance magnitudes at 1 kHz before and after coating. Error bars represent the standard deviation from the measurements. (K) Helium leak trace recording during testing of one array indicates that the sample is hermetic with no detectable leak.

## 2. RESULTS

### 2.1 Carbon Cybernetics Array Fabrication and its Hermeticity

Hermetic encapsulation remains a significant challenge for the development of high-resolution neural implants. Traditional encapsulation techniques have largely relied on titanium or ceramic housing, accompanied by ceramic feedthrough plates that contain an array of brazed wires. Increasing the electrode density poses a risk of the ceramics cracking, indicating the inherent limitation of this method. To overcome this constraint, we take advantage of diamond, an inherently impermeable material with extreme mechanical strength. Using gold active alloy brazing, we have pioneered and published a technique to achieve a hermetically sealed diamond encapsulation suitable for high-resolution neural implants^28^. Our Carbon Cybernetics array fabrication is based on this previous development.

The detailed manufacturing process for Carbon Cybernetics arrays is elaborated in the Methods section. Briefly, the fabrication begins by creating feedthrough vias and routing structures on the diamond substrate using laser milling. Individual carbon fibers are manually threaded into the feedthrough vias and secured with gold brazing. This is followed by mechanical polishing to remove the surplus braze and a cleaning step using reactive ion etching. Subsequently, the carbon fibers are cut into designed lengths with a laser and coated with a thin layer of Parylene-C. The electrode tips, once de-insulated using a flame torch or laser de-insulation method, are coated with PEDOT:PSS to boost their electrochemical properties.

We set our arrays at a fiber-to-fiber spacing of 100 µm. They have a density of 10,000 shanks/cm^2^ and are optimized with the fiber lengths of 1-2 mm, facilitating insertion into the rat cortex. Using laser cutting, we created an adjustable structure with fibers of different lengths (**Figure 1D**) to ensure simultaneous sampling of neurons both across a laminar plane and at varying cortical depths.

Individual carbon fibers were anchored in place using gold braze, serving the dual purpose of securing the fibers and establishing an electrical connection to the recording electronics. Such a connection could either be established via cables to an external device or integrated with a control ASIC for a fully implantable solution. **Figure 1B** shows an electric circuit built on the diamond surface using gold braze, tailored for a 5 × 5 carbon fiber array. This representation indicates the comprehensive coverage of the feedthrough holes, traces and connection pads by the gold braze, alongside the distinct isolation between electrode channels. The central 5 × 5 circular feedthrough structures on the diamond substrate were in direct contact with the carbon fibers. They were then connected to the larger rectangular pads near the edge of the diamond substrates. These larger pads, each measuring 100 µm × 50 µm, facilitated the wiring or ASIC integration.

Carbon fibers protruded from the diamond surface after threading. The extrusions were removed during the polishing process to clear away the excess gold braze. Following the polishing and reactive ion etching, some of the carbon fibers remained visible on the surface of the gold braze within the feedthrough holes, as shown in **Figure 1C, E**. Using Focused Ion Beam (FIB) technology, we assessed feedthrough cross-sections to examine the interaction between the carbon fibers and the braze. **Figure 1F** shows the close contact established between the carbon fibers and the braze. Ensuring a tight bond between the carbon fibers and the braze is critical as it directly determines both the conductivity and hermeticity of the array, which in turn further influences the efficiency and longevity of the device.

After applying an insulating and biocompatible layer of Parylene-C polymer, the carbon fiber electrode tips were exposed using flame torch or laser deinsulation. SEM measurements revealed that the lengths of the exposed fiber tips ranged between 40 µm and 200 µm. Uncoated carbon fibers had an average impedance of 385 kΩ at 1 kHz, which resulted in a high noise level for recordings. To reduce their electrochemical impedance, the carbon fibers were treated with a PEDOT:PSS coating. Successful application of this coating was verified via both SEM imaging and electrochemistry measurements (**Figure 1G-J**), which showed a reduction in impedance by 2 orders of magnitude, resulting in a value of 60 kΩ at 1 kHz.

After the array fabrication was completed, the hermeticity was demonstrated using helium leak detection tests. **Figure 1H** shows a typical helium leak detection trace recorded during the testing of one representative sample. Helium was introduced over the sample at the beginning of the test window. Large leaks result in an immediate sharp spike in helium detection, while small leaks (10^-9^-10^-7^ mbar L/s) result in a visible perturbation of the linear trace. An undisturbed line, such as the one depicted in Figure 1H, indicates that the helium leak rate is below the detection limit of the experiment (10^-11^ mbar L/s).

### 2.2 Chronic Biocompatibility and Safety of the Carbon Cybernetics Array

To assess the biocompatibility and safety of the Carbon Cybernetic array, we implanted the devices into the rat cortex for up to 6 months. In these experiments, the passive arrays, comprising 5 × 5 carbon fibers anchored to diamond substrates, were inserted without establishing any electrical connections or recordings. Following craniotomy and durotomy, the arrays were delicately inserted into the cortex of anaesthetized rats, facilitated either by vacuum tweezers or mechanically controlled tweezers (**Figure 2B**). For arrays with fiber lengths below 2 mm, insertion was seamless. **Figure 2C** shows the force measurements from an array with 25 fibers, each 2 mm long, during insertion and extraction in a rat brain. The array required only 1 mN to penetrate the tissue, with a peak insertion force about 2 mN as all fibers fully entered the tissue. Notably, we estimate that the force required to insert one carbon fiber is around 0.1 mN. Here, we see that this value escalates linearly with the number of fibers in an array. For chronic implantation, upon being released from the holding tweezers, the arrays remained securely positioned on the cortical surface (**Figure 2Bii**). We then sealed them in place using multiple layers of silicone and dental cement before animal recovery.

**Figure 2.**
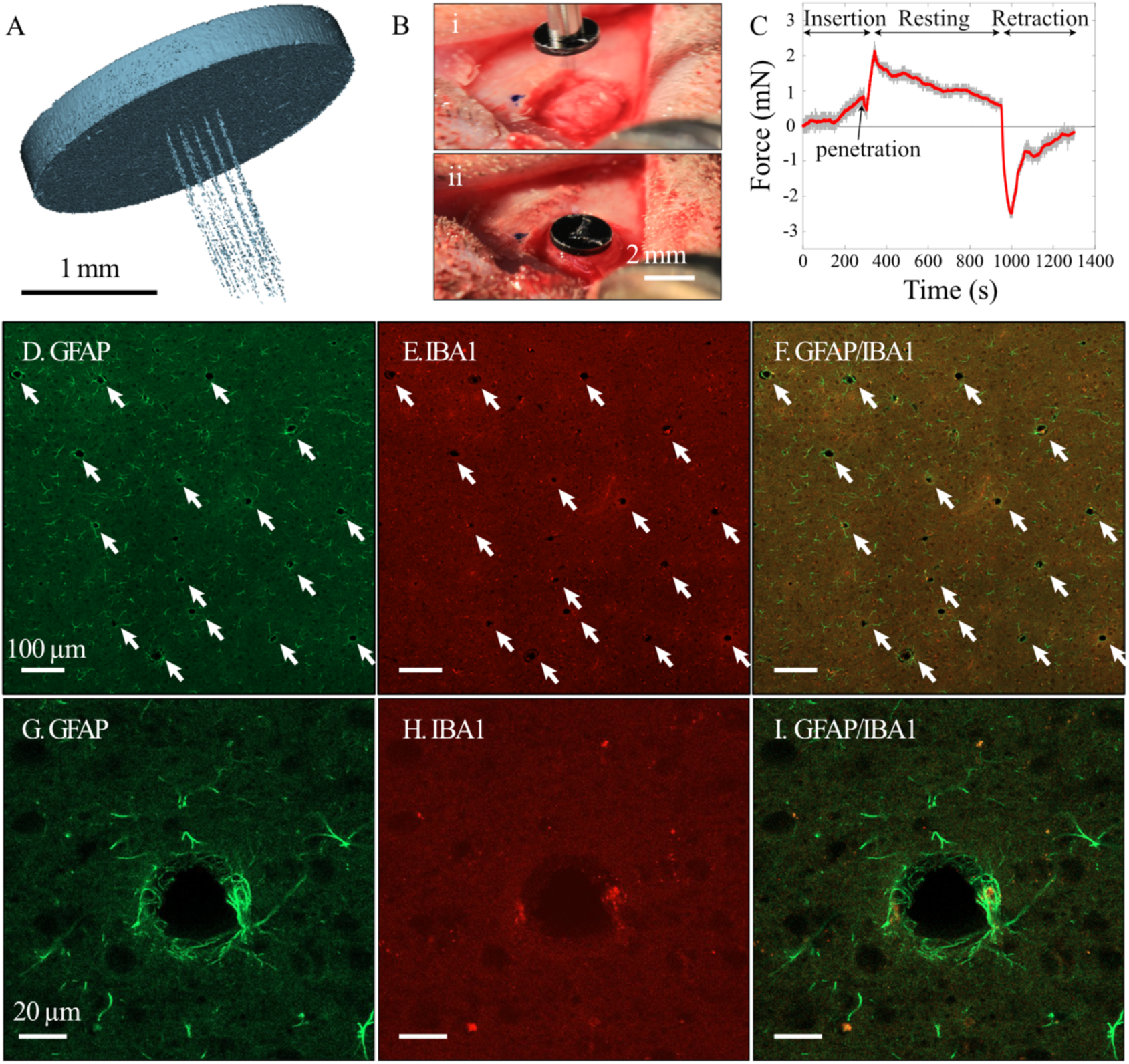
Implantation and Chronic Tissue Response of Carbon Cybernetic Arrays. (A) Micro-CT reconstruction of an array post six-month implantation in the rat cortex, showing intact fiber and secure attachment to the diamond substrate. (B) Visualization of successful implantation of an array with 25 carbon fibers, each 1-1.5 mm in length. (C) Force measurements during insertion and extraction of a 25-fiber array ( 2mm fiber length, insertion/extraction speed: 7 µm/s) in the rat brain. (D-I) Representative immunohistology of brain tissue post three-month implantation. Established glial scar markers, GFAP for astrocytes (D, G), and IBA1 for microglial (E, H) were used. F and I show combined views. Carbon fiber-induced holes are highlighted with arrows in D-F. Panels G-I show a close-up view of a representative hole. Overall, the immunohistology findings suggest minimal chronic tissue reactions to these carbon fibers. Scale Bar: D-F 100 µm, G-I 20 µm.

After the animals had been humanely killed, micro-CT scanning was performed, which provided insight into the positioning and condition of the implanted arrays. **Figure 2A** shows the micro-CT reconstruction of the array in an animal after 6 months of implantation. It confirms the intact and linear structure of the carbon fibers protruding from the diamond substrate, maintaining their originally designed tapered structure. **Supplement Video 1** shows the different layers of materials reconstructed from the scanning, showing the position of the array within the animal.

To evaluate the extent of glial scar formation, the arrays were explanted prior to histological analysis. We present representative fluorescent images of brain slices obtained after over 3 months of implantation **(Figure 2D-I)**. The histological evaluation highlighted a minimal tissue response elicited by the carbon fiber electrodes. Staining of astrocytes (using GFAP) and microglial cells (using IBA1), both major contributors to glial scarring, revealed only a thin scattering of these cells around the sites previously occupied by carbon fibers. We compared the extent of glial scars in animals from 3 months to 6 months of implantation (**Figure S1**). Notably, this layer thickness (<20 µm) was significantly reduced in comparison to glial scarring typically found with conventional neural probes (>100 µm)^9, 29^, regardless of the implantation duration. These collective findings underscore the minimally invasive nature of the carbon fibers and confirm the long-term biocompatibility and safety of the Carbon Cybernetics arrays.

### 2.3 Single-unit Recordings in Rats

To evaluate the capability of our Carbon Cybernetics arrays for recording, we wired the arrays consisting of 16 fibers with custom-designed flexible PCBs, which facilitated the connection to external recording systems (**Figure 3A**). We then implanted the arrays in the rat cortex and measured the spontaneous spiking activities of neurons while the animals were under anesthesia (**Figure 3B**).

**Figure 3.**
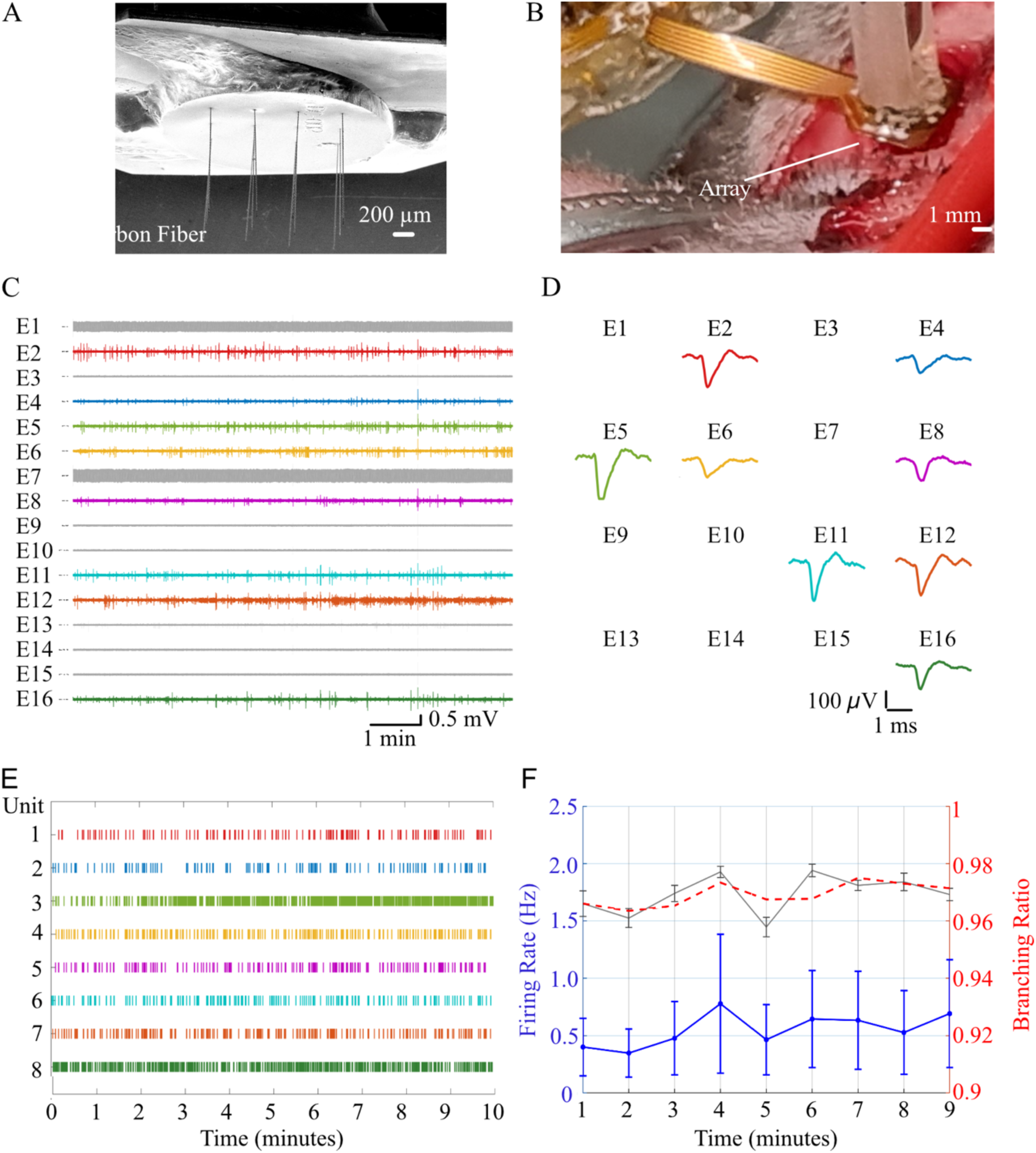
Acute Recording using Carbon Cybernetics Arrays in rat. The recordings were conducted using Carbon Cybernetic arrays with 16 electrodes connected to custom-designed PCBs. (A) shows an SEM image of an array. (B) The array was implanted in the rat cortex for recording while the animal was under anesthesia. (C). Recordings from a representative array, with extracted spike waveforms shown in (D) and spike train shown in (E). The firing rate (blue) and branching ratio (red and grey) are shown in (F). The grey line represents branching ratios calculated in 1-minute window, and the red dotted line represents the moving mean with a window of 5 minutes. The error bar shows the standard deviation.

**Figure 3C** presents a 10-minute recording from a representative array. In this array, 14 out of 16 channels exhibited low noise levels, typically ranging from 30 to 40 µV. Spike-sorting using Kilosort identified 8 channels that contained clear action potentials that could be extracted from the background noise (**Figure 3D**). For every single-unit, we extracted 10,000 random spike waveforms in the period between 1 ms before and 3 ms after the spike time.

The signal-to-noise ratio (SNR) was calculated for each single unit detected and was used to evaluate the quality of our recordings. SNR was determined by taking the ratio of the peak-to-peak amplitude of the mean waveform to the standard deviation of the background noise^30^. For the channels represented in **Figure 3D**, the SNRs varied between 1.9 and 4.0, with an average of 2.6. For a comprehensive analysis, we further classified the single units into different spike waveform classes (i.e. regular, fast, compound, triphasic, and positive spiking)^31^. The objective was to systematically determine if the same type of neurons were recorded from each electrode channel. The categorization of spike waveform classes was conducted in a semi-automated manner, using a waveform classification algorithm previously developed^31^. Notably, all the spikes recorded from our devices were classified as regular spiking, suggesting that the electrode recordings were made from excitatory neurons, most likely from pyramidal neurons in the cortex^32^.

**Figure 3E** shows the spike train extracted from the recording data shown in **Figure 3C** after spike detection. From the spike train, we quantified both the firing rate and the branching ratio (**Figure 3F**). The firing rate was between 0.25 and 0.75 Hz during the recording period, consistent with previous reports^33^ of spontaneous activity in animals under anaesthesia.

The branching ratio characterizes the propagation dynamics of activity within a neural network. It is defined as the ratio between the number of active neurons in one time step and those active in the preceding time step. A branching ratio close to 1 indicates that activity is self-sustaining, reflecting a critical state where excitation and inhibition are balanced. In contrast, a branching ratio below or above 1 suggests subcritical (fading) or supercritical (runaway) dynamics, respectively. During this recording, the branching ratio was between 0.96 and 0.98, consistent with previous studies^34^.

The capability of carbon fiber electrodes for long-term, chronic neural recording has previously been reported in several studies^20–23^. We also achieved stable single-unit recording in the rat cortex using carbon fiber electrodes encapsulated in a 3D printed design, with a consistent SNR over a period of three months (**Figure S2**).

### 2.4 Insertion and Single-unit Recordings in Sheep

To facilitate the future clinical translation of the arrays, we designed an insertion toolset for manually implanting the array via a small burr hole in the sheep’s brain. This insertion toolset enabled enabled controlled, precise and reproducible perpendicular placement of the electrodes into the cortex (**Figure 4A-C**). The system consists of three main components: a burr hole flange, a plunger, and a micrometer inserter. The burr hole flange is secured to the skull with stainless steel screws and serves as a guide to maintain perpendicular alignment of the electrode. The plunger, which holds the array, slides smoothly through the flange and connects magnetically to the micrometer via its top end. The micrometer inserter provides fine vertical control, allowing slow and steady penetration of the cortical surface while minimizing insertion force. The micrometer inserter can be removed after the successful insertion of the arrays, leaving the burr hole flange and plunger on the surface the skull for securing the arrays in chronic applications (**Figure 4C**). All components are easily assembled and fabricated using 3D printing technology with medical-grade MED-AMB 10 resin or stainless steel. Together, this system ensures accurate, stable, and minimally invasive implantation of the carbon fibre arrays. The successful insertion of the arrays using this toolset is evidenced via recording from sheep cortex, with one representative channel and the extracted single-unit shown in **Figure 4E,F**.

**Figure 4.**
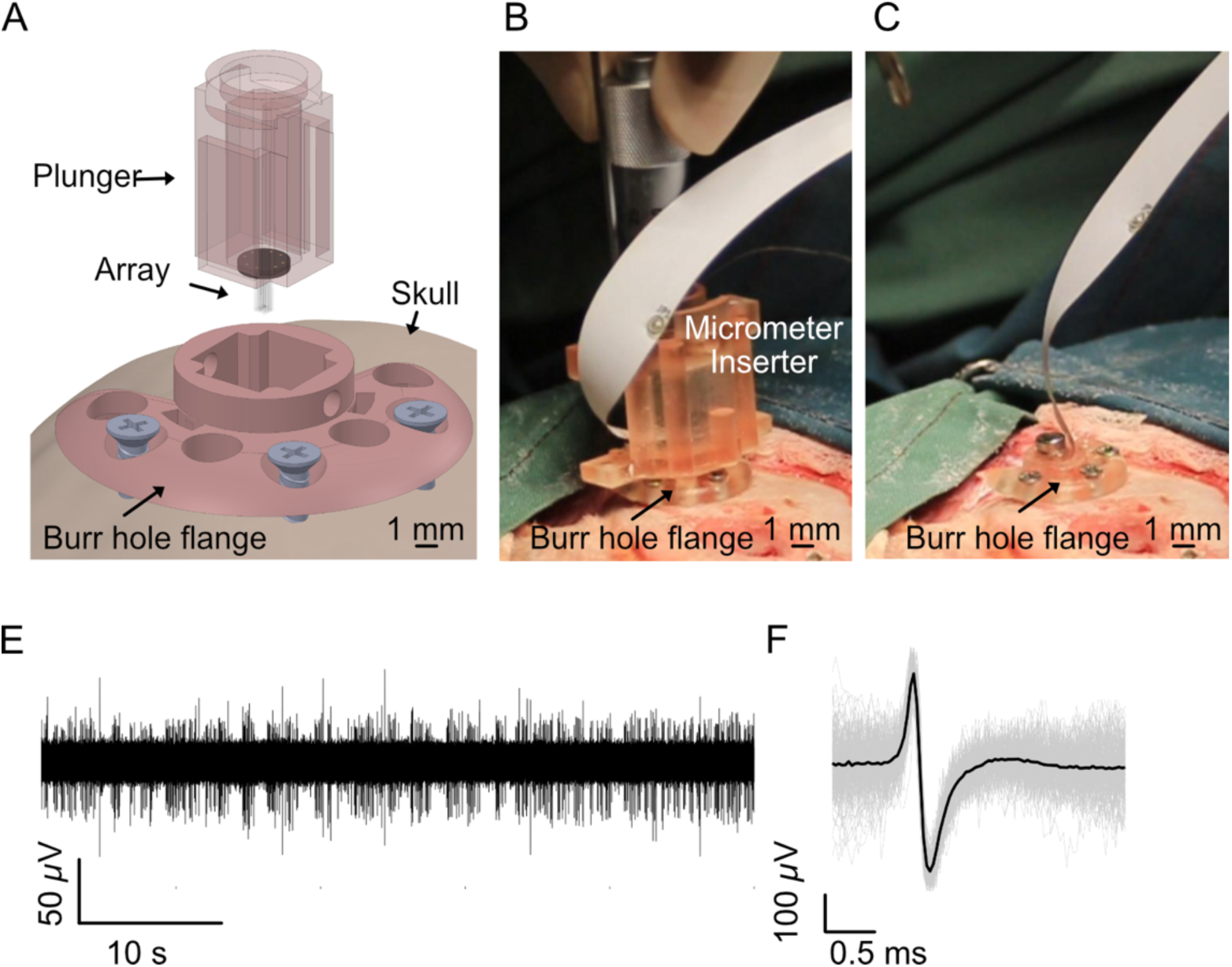
Manual insertion of Carbon Cybernetics Arrays in Sheep. (A) Schematic of the insertion system, including the plunger and burr hole flange. (B) Manual insertion of the array during surgery. (C). The array positioned on the cortical surface after removal of the micrometer inserter. (E) Representative recording from a single channel and (F) the corresponding spike waveform.

### 2.5 Scalability of Fabrication

A key advantage of our fabrication method was the laser milling process, which offered us the flexibility to modify the overall array configuration. Highlighting the scalability of our fabrication technique, our prototypes showcase arrays with channel counts from 25 electrodes (**Figure 1**), 100 electrodes (**Figure 5A**) and 1024 electrodes (**Figure 5B).** These arrays provide the potential for simultaneous neural recording of lateral information across different depths within a large volume of the brain. We envision the future development of a fully implantable device capable of recording from 1024 channels, optimized for lateral neural recording (**Figure 5C**). This approach would allow for comprehensive data acquisition while minimizing invasiveness, improving both the precision of the recordings and the long-term biocompatibility of the device.

**Figure 5.**
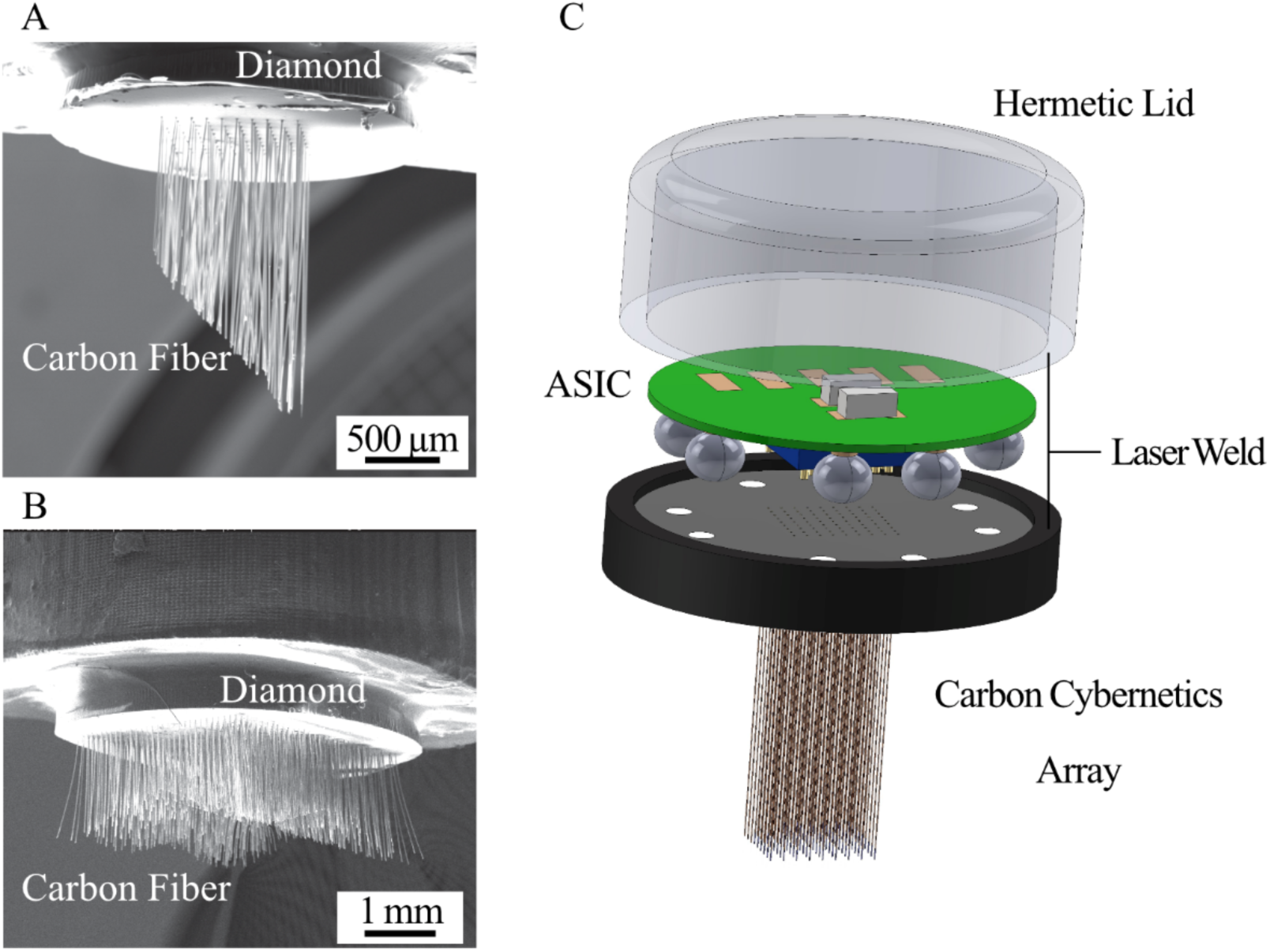
Versatile Configurations of Carbon Cybernetic Arrays and their Insertion. SEM images of: (A) 100 carbon fiber array and (B) 1024 fibers. All have a fixed fiber-to-fiber distance of 100 µm, reaching a channel density of 10,000 channels/cm^2^. Each configuration exhibits fibers trimmed to varying lengths, which allow for the simultaneous neural recording of lateral information across different depths of the brain. (C) A schematic representation of a compact, fully implantable Carbon Cybernetic Array device. The hermetic feedthrough design and the gold/diamond circuits of the Carbon Cybernetics Array will allow a direct interface with an Application Specific Integrated Circuit (ASIC) for data acquisition and wireless communication. This assembly will be encapsulated within a transparent, hermetic diamond lid, sealed by laser welding.

## 3. DISCUSSION

This study presents the development and assessment of a miniaturized, carbon-based microelectrode array for intracortical recording. Our unique use of diamond substrates, paired with the feedthrough design, addresses several key challenges in existing *in vivo* recording devices. Diamond is recognized as the most impervious, durable and biochemically inert material^35^. We had previously demonstrated the hermeticity and chronic safety of the feedthroughs constructed by diamond and gold active brazes^28, 36^. In the previous work, the electrode array was planar, with conductive diamond serving as the electrodes. This work provided further evidence of the hermeticity of the diamond and gold braze feedthrough structure (**Figure 1K**).

Building upon this foundation, we introduce a new array configuration that replaces the planar diamond electrodes with penetrating carbon fibers, further advancing the array’s capabilities for neural recording. Crucially, this hermetic feedthrough design paves the way for the integration of an encapsulated ASIC, as demonstrated in our previous work^27^. Additionally, we have developed a laser welding technique to create a hermetic seal between the array and a diamond lid, which can be engineered to remain transparent, enabling wireless communication via optics^28^. Together, these advances provide the groundwork for the development of a fully implantable, chronically stable device for medical and neuroscience applications (**Figure 4C**).

Our design employs laser micro-machining to define the feedthrough patterns. The flexibility of laser micro-machining ensures that the array’s configuration can be custom-tailored to specific application needs. In this work, we successfully fabricated an array with 1024 channels, achieving a density of 10,000 shanks/cm^2^ (**Figure 4B**). This equips us with the capability to record neural activity with high spatiotemporal resolution across millimeter-scale regions of the brain. By trimming the fibers into different lengths using the laser, we can simultaneously record across a large laminar plane but at different cortical depths. The potential for simultaneously recording across multiple different brain regions is crucial for neuroscience, offering insights into the complex neural network. Moreover, this expanded capacity is poised to enhance the potential of penetrating arrays for BMIs, offering higher degrees of freedom in motor or sensory transduction due to broader coverage^37^.

This work provides further evidence of the chronic safety of carbon fibers and their precision in neural recording. Consistent with prior studies^20, 21, 23, 26^, we observed minimal tissue responses evoked from the chronic implantation of carbon fibers (**Figure 2**). This is attributed to the small dimension and the inherent flexibility of the carbon fibers. The small dimension also allows for the recording of single units, as demonstrated in **Figures 3, 4**. Both characteristics will ensure the use of these devices for robust, long-term measures of neural activity, in research settings and clinical applications.

The design of Carbon Cybernetics arrays also facilitates the option of suturing the dura closed after implantation, similar to the Utah array. In Neuralink’s first clinical trial, there were reports of flexible threads inserted into the cortex being pushed out of the brain^8^. This may be attributed to the challenge of closing the dura after implanting the threads, as scar tissue formation at the entry point could cause the threads to withdraw. In contrast, the ability to suture the dura after implantation, as is possible with the Utah array design, may minimize this issue and therefore is advantageous for long-term stability.

Here, we also designed a low-cost, 3D-printed insertion toolset that enables manual yet reproducible implantation of the arrays into the sheep cortex. This toolset can be readily adapted for use in human surgery and allows electrode insertion through a small burr hole. The design eliminates the need for costly surgical robotics and avoids the risks associated with large craniotomies, thereby enhancing the translational potential of the Carbon Cybernetics arrays.

The promise of the Carbon Cybernetics arrays in neuroscience is vast, offering avenues to decipher neuronal circuits, brain development, behavior and consciousness^38^. At the therapeutic level, precise and chronic brain signal monitoring can reveal the patterns of neurological and psychiatric diseases, and provide insights into disease prediction and prevention, leading to innovative therapies^17^. Therefore, our envisioned ASIC-integrated, fully implantable Carbon Cybernetics arrays, with their long-term reliability and safety, are poised to bring significant benefits to the advancement of the treatments for these diseases.

While we have achieved the fabrication of arrays with 1024 fibers, it has yet to be tested for insertion and neural recording. Wiring up such high channel arrays to external devices or control ASICs remains a challenge that requires novel solutions but is out of the scope of this study. In our tests, 5 × 5 arrays with fibers exceeding 2 mm in length posed insertion challenges, hence we capped fiber lengths at 2 mm. However, recording from deeper brain areas, such as the hippocampal region, offers considerable value in many applications. To facilitate the insertion of longer fibers, temporary toughening of the fibers with dissolvable chemicals, as shown in our prior work^39^, may be helpful. Future work will also include automation of carbon fiber assembly through robotic integration, improving both product yield and quality consistency.

## 4. METHODS

### 4.1 Array Fabrication

Feedthrough structures in the diamond substrate were created as previously described^29, 36^. Briefly, polycrystalline diamond wafers (Element Six Ltd) were polished using a Coborn PL3 rotary polisher. They were then laser-cut into 2.5 mm diameter circles using an Oxford Lasers Alpha series with an Nd: YAG Laser at 532 nm. Feedthrough holes, set at a 100 µm pitch, were fabricated into the substrate through laser milling. This method was also employed to establish tracks for wire routing and contacting pads. To eliminate graphitic residues, substrates were boiled in a 10 ml H_2_SO_4_ solution with 1 g NaNO_3_ for 60 minutes.

Following acid boiling, 200 nm layers of both Mo and Nb layers were sputter deposited on the prepared diamond surface using an RF/DC Anatech Hummer. Single carbon fibers were manually threaded into the feedthrough holes, protruding from the diamond surface. To anchor these fibers and form necessary conductive tracks, Au-ABA braze paste was applied on the diamond and annealed at 1100 °C for 30 minutes. Excess braze was polished off, and residue was removed through reactive ion etching^40^. The fibers were then laser-trimmed to lengths between 1 and 2 mm. Finally, the carbon fiber side was coated with 2 µm of Parylene-C using a PDS 2010 system (SCS, Specialty Coating Systems).

To build an electrical connection between the array and the external recording system, we used a custom-designed flexible PCB. Silver paste was used to forge a connection between the PCB and the gold pads on the diamond substrate. Connected joints were subsequently encapsulated in epoxy.

Carbon fiber electrodes were exposed and deinsulated using a blowtorch, while immersed in deionized (DI) water. Electrode impedance was reduced by electrochemically depositing a layer of PEDOT:PSS. Cyclic voltammetry, in a range of -0.5 V to 0.9 V at a rate of 100 mV/s, was conducted for 15 cycles in a PEDOT:PSS (1:2) solution using the Gamry Interface E1000. Successful deposition was verified through electrochemical impedance spectroscopy (EIS) measurements and SEM imaging.

### 4.2 Hermeticity Testing

Hermeticity tests were conducted using an Adixen ASM310 Portable Helium Leak Detector on the arrays. Samples were mounted into a custom-made holder such that the arrays were clamped between two Viton O-rings. One side of the sample was exposed to the atmosphere, and the other side to the vacuum of the helium leak detector. Once baseline vacuum pressure was achieved, a slow flow of helium was introduced to the array. The detector has a detection limit of 10^-11^ mbar L/s.

### 4.3 Surgical Procedures

#### Rat Experiments

All rat experiments were conducted in accordance with the Australian Code for the Care and Use of Animals for Scientific Purposes (National Health and Medical Research Council, Australia) and approved by the University of Melbourne Animal Ethics Committee (Ethics Approval #20180 and #22494).

Adult Long Evans rats of either sex (sourced from Ozegene ARC, Australia) were used for electrophysiological recordings. Anaesthesia was induced with 5% isoflurane and maintained at 1.5% using a SomnoSuite system (Kent Scientific). Body temperature was maintained using a feedback-controlled heating pad. A 6 × 6 mm craniotomy and duratomy were performed over the primary visual cortex (V1). Stainless steel screws were secured to the skull to provide electrical grounding, and the cortical surface was kept moist with sterile saline-saturated gel foam. The dura mater was carefully incised under a surgical microscope using fine microsurgical scissors or a 30-gauge needle to minimize trauma and avoid major surface vessels. A small dural flap (∼1–2 mm²) was retracted to expose the underlying cortex. The exposed region was continuously hydrated with sterile saline.

Carbon fiber microelectrode arrays were positioned above the target site using a mechanically controlled tweezer and micromanipulator (Scientifica, UK) and inserted vertically into the cortex to depths of 500–1500 µm. After insertion, the array was stabilized with surgical silicone adhesive (Kwik-Sil, World Precision Instruments) and encapsulated with UV-curable dental cement (3M). Following surgery, animals received postoperative analgesia consisting of Buprenorphine and Meloxicam for three consecutive days and were monitored twice daily for two weeks. At the conclusion of experiments, animals were humanely euthanized via intracardiac injection of sodium pentobarbital, followed by transcardial perfusion with 4% formalin. Brains were stored at 4 °C prior to histological analysis.

#### Sheep Experiments

Experiments were conducted at the Old Howard Florey Institute and approved by the Florey Animal Ethics Committee (Approval ID: 22-010-UM).

Experiments involving sheep were conducted under both local and general anaesthesia to ensure complete analgesia and muscle relaxation. Anaesthesia was induced via intravenous injection of sodium thiopentone (5 mg/kg) and maintained using 1.5–4% isoflurane in oxygen, adjusted according to body size. Heated air delivered through the endotracheal tube maintained body temperature between 36 °C and 38 °C. Temperature, respiration rate, and anaesthetic depth were monitored every 30 min throughout the procedure. The head was shaved and disinfected three times with Betadine, and ophthalmic ointment was applied to protect the cornea. The animal was then secured in a stereotaxic frame. A midline incision was made along the scalp, and the periosteum was removed to expose the skull surface. A 10–14 mm burr hole was drilled to expose the visual cortex, and the dura mater was carefully incised under direct visualization to expose the cortical surface. The dural window was opened sufficiently to allow electrode access while minimizing cortical disturbance.

Electrodes were inserted into the cortex to a depth of up to 2 mm using either custom-designed insertion tools or a micromanipulator. The exposed cortex was continuously irrigated with sterile saline to prevent desiccation. Haemostasis was achieved through gentle cauterization or compression using sterile gauze when required.

At the conclusion of the experiments, animals were deeply anaesthetized with an overdose of sodium pentobarbital administered intravenously until cardiac and respiratory activity ceased. Transcardial perfusion was then performed with phosphate-buffered saline followed by 4% formalin to ensure fixation of the brain. Brains were carefully extracted and post-fixed in 4% formalin at 4 °C for subsequent histological and structural analysis.

### 4. Insertion Tool

To ensure precise, reproducible implantation of carbon fiber arrays into the cortex, a set of device insertion accessories was used to control and guide insertion. These components facilitate slow, perpendicular penetration of the cortical surface while minimizing insertion force. All accessories are easily assembled, affixed to the skull using standard screws, and fabricated from medical-grade MED-AMB 10 resin and stainless steel (Figure 4).

#### Burr Hole Flange

The burr hole flange is secured to the skull using three standard self-tapping stainless steel screws (2G × 6.5 mm). Once affixed, the flange functions as a guide for the plunger, ensuring perpendicular alignment of the electrode relative to the cortical surface.

#### Plunger

The diamond electrode array is attached to the bottom of the plunger, which slides through the burr hole flange. Each side of the plunger includes a thin channel to accommodate the PCB without obstructing vertical movement. A magnet located at the top of the plunger interfaces with the micrometer to allow controlled advancement.

#### Micrometer Inserter

The micrometer inserter is mounted atop the burr hole flange and provides alignment and support for the micrometer. The micrometer is positioned through the top portion of the inserter and magnetically connected to the plunger. Using the micrometer, the operator can precisely control the insertion speed of the electrode array into the cortex.

All components are designed to maintain perpendicular orientation and smooth vertical movement during implantation, ensuring accurate and minimally invasive electrode placement.

### 4.4 Insertion Force Measurement

The insertion and extraction forces of the Carbon Cybernetics Arrays were measured using a Mark-10 Series 7 Force Gauge, sampling at approximately 21 Hz. Both insertion and extraction speeds were maintained at 7 µm/s.

### 4.5 Electrophysiology Recordings and Data Analysis

Extracellular recordings were acquired at 30 kHz using a Ripple acquisition system (Ripple Neuro) and visualized using Trellis software. The raw data were high-pass filtered at 300 Hz to remove low-frequency noise. To separate the spikes from different neurons, we used an automatic spike-sorting program called Kilosort^41^, and spike clusters were further manually curated using the graphical user interface Phy^42^.

We collected 10,000 random spike waveforms for each single unit from the raw data. These waveforms were extracted in the period between 1 ms before and 2 ms after the spike time (in a window of -30 to 60 samples, where 0 is the minimum trough). We used the maximum number of spikes for single units with less than 10,000 spikes. Collected spike waveforms were averaged, and the baseline was subtracted to make the new baseline equal to 0. The baseline was calculated as the mean of the first and last ten samples of the extracted waveform. The signal-to-noise ratio (SNR) was calculated as the ratio between the mean waveform and the standard deviation of waveform noise, as described by Kelly *et al.*^30^. An unpaired two-sampled t-test was used to determine the statistical significance of the differences in SNRs between different time points.

Only well-isolated single units with refractory periods ≥1.5 ms were included in the following Firing rates and Branching ratios analyses. For each unit, the mean firing rate was computed as the total number of spikes divided by the recording duration. Across units, the average spontaneous firing rate under mild isoflurane was >1 Hz.

To quantify the propagation of activity across time, we estimated the branching ratio (σ), defined as the expected number of descendant events triggered by one preceding event. We applied the multistep regression (MR) estimator following Priesemann *et al.*^34^. Spike trains from all single units were binned into time steps of width Δt (chosen to match the mean inter-event interval, typically 2–4 ms). Denoting the total number of active units (or spikes) in bin t as A_t_, the MR estimator uses the collection of linear regression slopes:

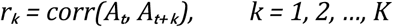

measured over multiple time lags k. Under the assumption that r_k_ ∝ σᵏ (which holds even under subsampling), the branching ratio is then estimated as:

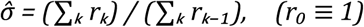

This approach corrects for subsampling and temporal correlation biases, providing a robust estimate of the true underlying branching process. Analyses were implemented in MATLAB using the publicly available MR-estimator toolbox^34^. To assess robustness, we also computed the conventional direct-ratio estimator; although absolute values differed, the same trends emerged.

### 4.6 MicroCT Scanning and Image Reconstruction

To confirm the location of the array after long-term implantation, a micro-CT scan was conducted with the perfused rat head. 4% formalin was used to fixate the rat head for 24 hours. The head was scanned at low resolution (10.3 µm) to image the full head and at high resolution (3 µm) to image the array and the fibers inside the brain tissue. The segmentation of micro-CT images was conducted using the open-source image analysis software 3D-Slicer (Version 5.2.2)^43^. The fiber elements extracted from the high-resolution images were further re-engineered to enhance their spatial definition using Ansys SpaceClaim (Version 2021R2). Following this, the 3D models reconstructed from both the low- and high-resolution datasets were combined to form an integrated model. This model accurately depicts the spatial interrelation of the fibers, the substrate, and the skull, providing a detailed representation of the structural configuration.

### 4.7 Immunohistology and Data Analysis

The implants were removed from the brain tissues before immunohistology. This was performed after skull decalcification by incubating the samples in 0.25 M EDTA solution at 4 °C for at least a week, following the protocol described in ref^26^. The brain sample was then prepared as brain slices or cleared using a tissue clearing kit (ab243298, Abcam) according to the manufacturer’s instructions. Primary antibodies used for immunohistology were: GFAP in chicken (ab4674, Abcam) and IBA1 in rabbit (Fujifilm Wako). Secondary antibodies were goat anti-chicken Alexa Flour 488 (ab150169, Abcam) and goat anti-rabbit Alexa Flour 647 (ab150083, Abcam). The samples were imaged using a Z-stack scan on either a multiphoton microscope or a light sheet microscope. The quantification of GFAP and IBA1 was conducted using ImageJ software^44^, and the location of the electrodes was manually determined.

## Data Availability Statement

All data needed to evaluate the conclusions in the paper are present in the paper and/or Supplementary Materials. Additional data related to this paper may be requested from the authors.

## Code Availability Statement

Kilosort2 was used for the automatic sorting of spikes and is available at https://github.com/cortex-lab/KiloSort. A graphical user interface, *Phy* was used for manual clustering of automatically sorted spikes and is available at https://github.com/cortex-lab/phy.

## Acknowledgements

The authors acknowledge the facilities, and the scientific and technical assistance of the RMIT Microscopy & Microanalysis Facility (RMMF), a linked laboratory of Microscopy Australia, enabled by NCRIS. The authors acknowledge the staff at the Biological Optical Microscopy Platform (BOMP), the Melbourne Advanced Microscopy Facility, for providing technical assistance with tissue scanning and analysis. WT is supported by the Australian Research Council via a Discovery Early Career Researcher Award (DE220100302) and an ARC Linkage Grant (LP180100638). HW is supported by the Research Training Program Scholarship and a GCI top-up Scholarship. YJJ is supported by the Lions Vision Research Fellowship. DJG is supported by an ARC Future Fellowship (FT190100215). Carbon Cybernetics is supported by an MRFF, MTPConnect via Biomedtech Horizons (BMTH) grant (3_15). This research is also supported by a grant from the Australian College of Optometry.

