## Supplementary Materials for "Carbon Cybernetics Array: a miniaturized carbon-based microelectrode array for intracortical recording"

*Simon Higham et al.*

**This file includes:**

- Figure S1: Detailed analysis of tissue response to the Carbon Cybernetics arrays
- Figure S2: Long-term recording stability in rat cortex using carbon fiber electrodes

**Other Supplementary Materials for this manuscript include the following:**

- Supplementary Video S1: MicroCT reconstruction of a Carbon Cybernetics array after six months of implantation in the rat brain.

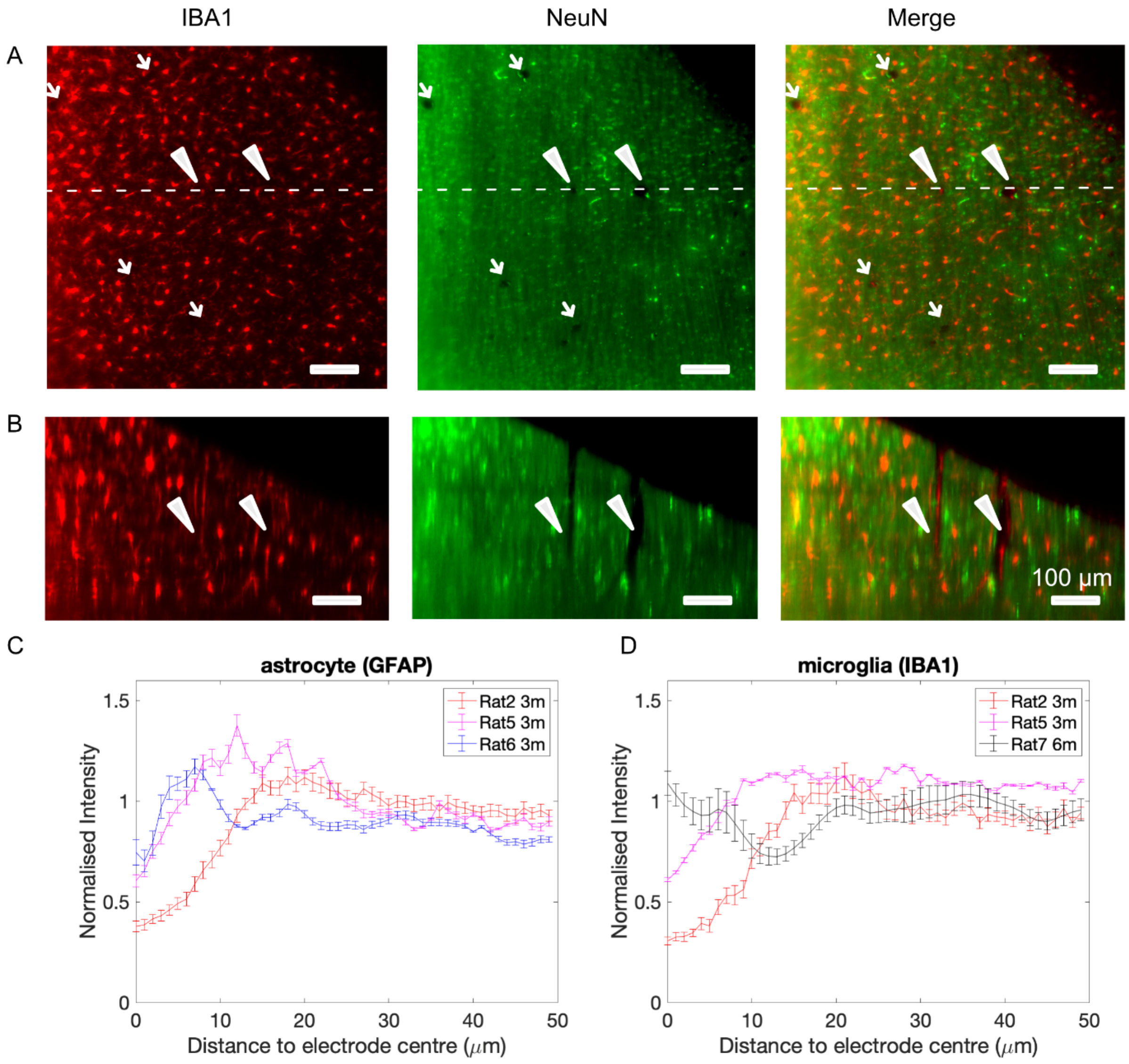


**Figure S1. Detailed Analysis of Tissue Response to Carbon Cybernetics Arrays.** (A). Fluorescent imaging depicting microglial (IBA1) and neuronal (NeuN) staining in brain tissues six months post-array implantation. Notably, arrows and triangles highlight the damage caused by carbon fiber electrodes. (B) Orthogonal view of depth images with a dashed line marking the perspective origin in (A). Scale bar: 100 µm. (C) and (D) show the normalized fluorescence intensity of GFAP and IBA1 profile as a function of distance from the center of the electrode tract, which is set as x=0. Some samples exhibited unsuccessful

GFAP or IBA1 staining and, therefore, were excluded from the plots. Error bars indicate the Standard Error of the Mean (SEM).

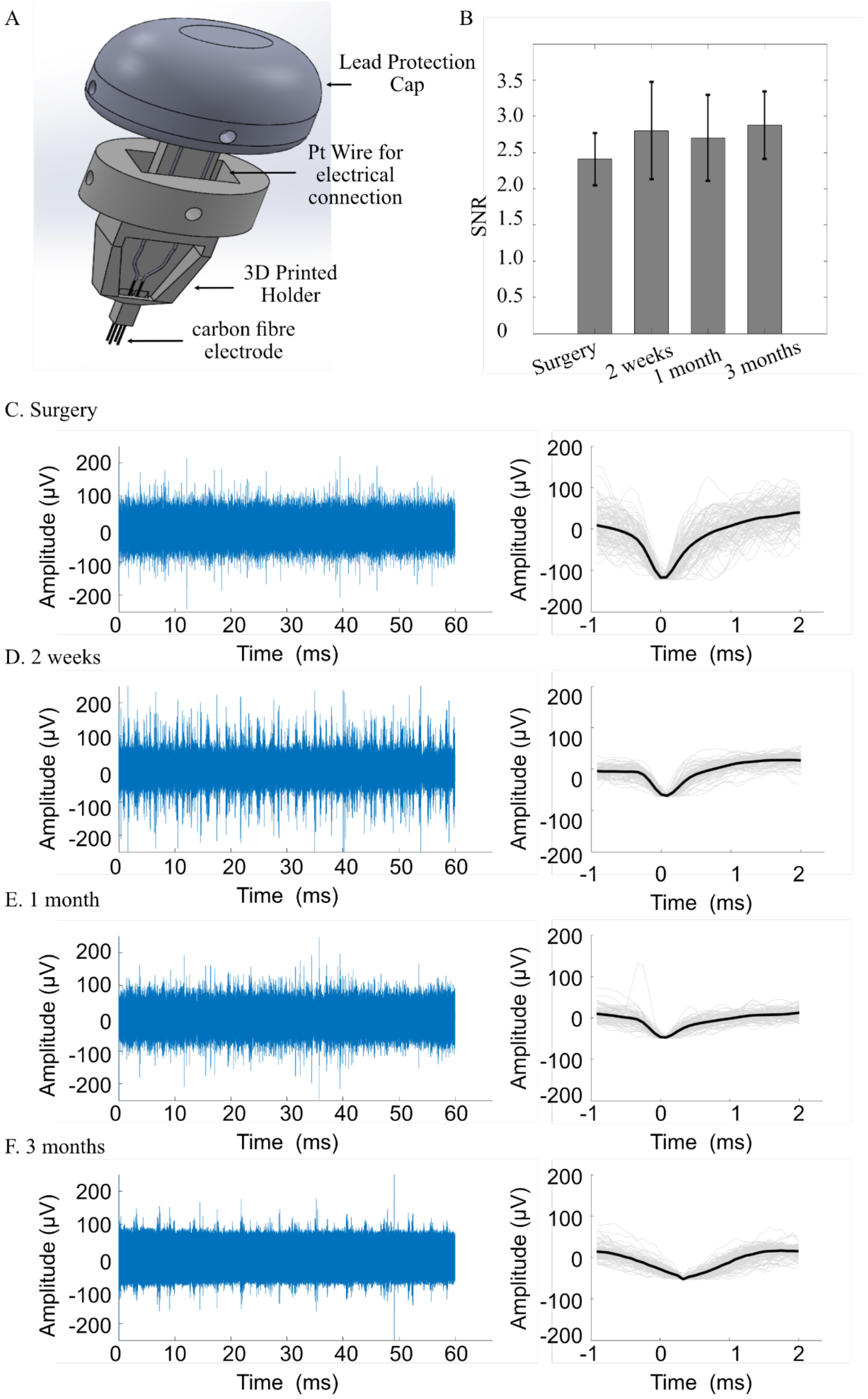


**Figure S2. Long-term Recording in Rat Cortex Using Carbon Fiber Electrodes.** A. Chronic recordings were achieved with carbon fibers secured in a custom 3D printed holder. These electrodes were connected to external devices via platinum wires, with a cap (also 3D printed)

protecting the connections. B. Recordings were performed intermittently: during surgery and then at 2 weeks, 1 month and 3 months post-surgery. All recordings were conducted while the animals were under anesthesia. The signal-to-noise ratio (SNR) of a representative electrode is depicted, illustrating stable recording quality over time. The error bar represents the standard deviation. We found no significant difference in SNRs between different time points using an unpaired two-sampled t-test. C-F show the raw signals (left panel) and the spike-sorted waveforms (right panel) from the representative electrode in B at different time points.
